# Histone lactylation H3K18la and acetylation H3K27ac encode two separable epigenomic features for enhancer regulation

**DOI:** 10.64898/2026.04.14.718598

**Authors:** Tetsuro Komatsu, Takeshi Inagaki

## Abstract

Histone lactylation H3K18la marks tissue-specific enhancers in combination with H3K27ac. It remains unclear whether and how these modifications differ. In this study, we dissected the specific roles of H3K18la- and H3K27ac-marked regions. We found that H3K18la predominantly localized at non-promoter regions with lineage-specific transcription factors and was strongly associated with enhancer activity and p300/CBP dependency. In contrast, H3K27ac was largely independent of these features; however, H3K27ac-occupied regions were selectively bound by the super-enhancer-enriched transcriptional coactivators BRD4 and MED1 and were responsible for gene expression involving liquid–liquid phase separation. The genomic regions covered by both H3K18la and H3K27ac best overlapped with super-enhancers. Thus, our study suggests that p300/CBP-dependent enhancer potential and BRD4/MED1-mediated enhancer cooperativity are separately encoded by H3K18la and H3K27ac, respectively, and exert synergistic enhancer activation when combined. Although the characteristics of H3K18la were similar to those of N-terminal acetylation of H2B (H2BNTac), a recently described enhancer signature, we found distinct genomic distributions of these marks. Collectively, our findings will advance the current understanding of how histone modifications establish enhancer landscapes.

## INTRODUCTION

Post-translational modifications of histones play a major role in epigenetic regulation by dynamically orchestrating genomic functions and chromatin structure within the nucleus. Therefore, the “histone code”^1^, namely combinations of individual modifications and their specific functions, has been extensively analyzed by researchers. For instance, mono- and tri-methylation at lysine 4 of histone H3 (H3K4me1 and H3K4me3, respectively) are known to localize to enhancers and active promoters, respectively, and serve as functional markers for these genomic elements^2^. Similarly, acetylation of histone H3 at lysine 27 (H3K27ac) has been widely used as a marker for active enhancers, ever since the report that this modification can discriminate active from poised enhancers^3^. However, the same study also revealed that a significant number of promoters are H3K27ac-positive across several cell types^3^, making it difficult to utilize this histone modification as the sole marker for active enhancers. Recently, it has been reported that acetylation of multiple lysine residues at the N-terminus of histone H2B (H2BNTac) co-localizes with H3K27ac at active enhancers; however, this modification poorly marks constitutively active promoters in mouse embryonic stem cells (mESCs)^4^. Thus, these findings establish H2BNTac as a more suitable marker for active enhancers. Both H3K27ac and H2BNTac can be catalyzed by the histone acetyltransferase p300 and its paralog CBP (hereafter p300/CBP), which are the central regulators of enhancer-dependent transcription^5^. H2BNTac outperformed H3K27ac in predicting p300/CBP-target genes/genomic regions^4^, suggesting that p300/CBP-dependent regulation may not be associated with H3K27ac. Indeed, despite the critical roles of p300/CBP at enhancers^6^, a study on mESCs harboring mutated histone H3.3 (K27R) suggested that a decrease in H3K27ac levels at enhancer regions results only in limited effects on the transcriptome^7^. In summary, a growing body of evidence contradicts the widely accepted notion that the presence of H3K27ac can be considered as a definitive indication of active enhancers.

Super-enhancers play a prominent role in the control of cell identity and possess strong enrichment of lineage-specific transcription factors (TFs), as well as transcriptional coactivators and chromatin regulators, such as p300, BRD4, and Mediator and cohesin complexes^8-10^. BRD4 and the Mediator subunit MED1 form nuclear droplet-like condensates through liquid–liquid phase separation (LLPS), gathering neighboring enhancers into a super-enhancer^11^. Although super-enhancers are generally considered to be clusters of typical enhancers in close genomic proximity^12^, a comprehensive characterization of the endogenous alpha-globin super-enhancer suggests the presence of “facilitators” as another constituent regulatory element^13^. Facilitators exhibit no intrinsic enhancer activity; however, they are required for the full potential of typical enhancers within the same super-enhancer. Thus, super-enhancer activities may be defined by the strength of individual constituent enhancers and by inter-enhancer cooperativity, which is mediated by facilitators and/or by the nature of BRD4 and MED1 to form liquid droplets.

Histone lactylation is a new class of modifications that originates from lactate, a metabolite mainly generated through glycolysis^14^. It has been demonstrated that changes in intracellular lactate levels—for example, upon inhibition of lactate dehydrogenase and under hypoxia—modulate histone lactylation levels, leading to epigenomic remodeling. Therefore, histone lactylation reflects cellular metabolic states in epigenetic and transcriptional regulation. *In vitro* biochemical assays using reconstituted chromatin templates have revealed that p300 catalyzes histone lactylation, which *per se* promotes transcription. The role of p300 in histone lactylation in cells was verified by the overexpression and knockout of this enzyme. Thus, p300 may play a role in epigenetic regulation through differential or cooperative actions on histone acetylation and lactylation.

Similar to acetylation, lactylation occurs at multiple lysine residues in all core histones^14^, and histone H3 lysine 18 is one of the best-studied lactylation sites (H3K18la). Recently, high-quality Cleavage Under Targets and Tagmentation (CUT&Tag) datasets for H3K18la were generated using multiple tissues and cell types^15^, showing that H3K18la is distributed on the genome in a manner resembling that of H3K27ac, with enrichment at active promoters and tissue/cell type–specific enhancers. However, the degree of (dis)similarities between H3K18la and H3K27ac has not been well-documented, leaving the specific functions of each modification, if any, largely unknown. Therefore, to examine whether and how H3K18la exhibits unique distribution on the genome, we conducted a comparative analysis of histone modification profiles using a wide range of public omics datasets (Table S1), including the aforementioned H3K18la CUT&Tag data.

## RESULTS

### H3K18la and H3K27ac jointly and individually localize on the genome in adipocytes

White adipocytes are metabolically active cells in which lactate production occurs dynamically in response to cues from the intra-/extracellular environment^16,17^. Our previous studies have elucidated the crucial roles of metabolic–epigenetic crosstalk in adipocyte fate determination^18,19^. Therefore, we first analyzed the H3K18la CUT&Tag datasets together with those for other modifications in mature adipocytes *in vivo*^15,20^. Focusing on the Assay for Transposase-Accessible Chromatin using sequencing (ATAC-seq) peaks in adipocytes^21^ (∼74k peaks), H3K18la and H3K27ac levels were examined for clustering to divide the peak regions into two groups, High and Low, for each modification (Figure S1A). Then, all ATAC-seq peaks were re-clustered into four groups covering all possible combinations (Figure 1A): H3K18la-High/H3K27ac-High (Cluster A1), H3K18la-High/H3K27ac-Low (A2), H3K18la-Low/H3K27ac-High (A3), and H3K18la-Low/H3K27ac-Low (A4). Consistent with the original study that reported similar distributions of H3K18la and H3K27ac^15^, we observed that a significant number of ATAC-seq peaks were marked by both modifications (Cluster A1, 9,812 regions). In addition, we found genomic regions that were occupied by either H3K18la (A2, 1,712 regions) or H3K27ac (A3, 5,731 regions), suggesting the presence of functions specific to each modification. To characterize the genomic loci singly or dually marked by H3K18la and H3K27ac, we analyzed H3K4me3, H3K4me1, and H3K27me3 levels in these regions as markers of active promoters, poised/active enhancers, and repressive domains, respectively (Figure 1A). As a global trend, H3K18la-marked regions, irrespective of H3K27ac co-occupancy, were found to be more strongly associated with H3K4me1 (Clusters A1 and A2), whereas those occupied only by H3K27ac were co-localized with H3K4me3 rather than with H3K4me1 (Cluster A3). H3K27me3 was not enriched in Clusters A1–A3, validating H3K18la and H3K27ac as active histone modifications. Consistent with the correlation with H3K4me3 and H3K4me1, we found that both Clusters A1 and A2 were mainly located in intergenic and intronic regions (∼60–70%); however, the majority of the regions in Cluster A3 resided in promoters (>70%, Figure 1B). These results suggest that H3K18la and H3K27ac have not only shared, but also unique genomic distributions, with the former being more associated with characteristics of enhancers in mature adipocytes.

**Figure 1.**
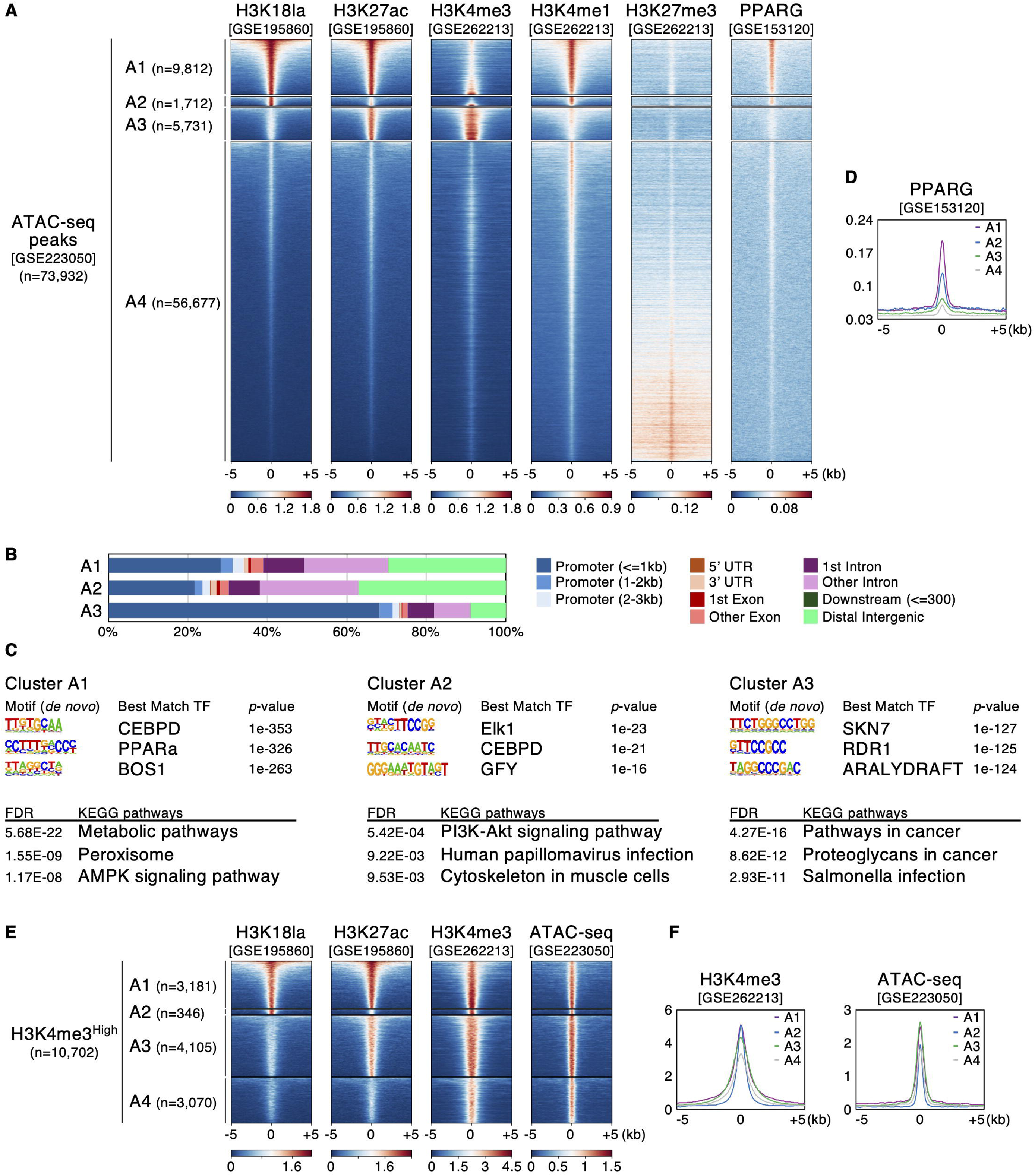
Cell type–specific genomic features are more associated with H3K18la than H3K27ac in mature adipocytes. (A) Heatmaps for CUT&Tag and ChIP-seq signals of the indicated modifications and factors at ATAC-seq peak clusters based on H3K18la and H3K27ac levels in mature adipocytes (peak center -/+ 5kb). Number of regions in each cluster are indicated on left. (B) Genomic distributions of peaks in Clusters A1–A3. (C) TF motif (upper panels, *de novo* motifs) and KEGG pathway analyses (lower panels, with peak annotation) for Clusters A1–A3. Top 3 hits for each analysis are listed. (D) Aggregation plot for PPARG ChIP-seq signals at each cluster. (E) Heatmap visualization of signals for the indicated data at H3K4me3-High regions of each cluster. (F) Aggregation plots for H3K4me CUT&Tag and ATAC-seq signals at H3K4me3-High regions of each cluster.

### H3K18la-marked regions represent adipocyte-specific genomic features more specifically than H3K27ac-marked ones

We performed motif analysis of TF recognition sequences for Clusters A1–A3 to examine whether the binding of specific TFs correlated with H3K18la or H3K27ac localization (Figure 1C, upper panels, Table S2). The top hit motifs in Cluster A1 included those for the CCAAT/enhancer-binding protein (CEBP) and peroxisome proliferator-activated receptor (PPAR) families. CEBPA and PPARG, which are master regulator TFs in adipocyte differentiation^22,23^, belong to these families. Similarly, the motif for CEBP TFs was enriched in Cluster A2. In contrast, Cluster A3 showed no enrichment of CEBP or PPAR recognition sequences as the top hits. These results led us to test whether the actual binding of adipocyte-specific TFs also exhibited cluster-specific distributions. To this end, we analyzed public Chromatin Immunoprecipitation followed by sequencing (ChIP-seq) data for PPARG in mature adipocytes^24^; to our knowledge, no studies have generated CEBPA ChIP-seq profiles using *in vivo* adipocyte samples (Figures 1A and 1D). Consistent with the motif analysis, Cluster A1 was most bound by PPARG. Although PPARG-related sequences were not detected in the motif analysis, we observed its localization in Cluster A2, which is in line with a previous study reporting that CEBPA recognition motifs are present in the vicinity of most PPARG-binding sites on a genome-wide scale^25^.

We also conducted a Kyoto Encyclopedia of Genes and Genomes (KEGG) pathway analysis for each cluster after peak annotation of the nearest genes (Figure 1C, lower panels, Table S3). The top hit in Cluster A1 was Metabolic pathways, which are central to adipocyte function. In the results for Cluster A2, the top hit was the PI3K–Akt signaling pathway, which plays a pivotal role in insulin signaling critical for adipocyte regulation^26^. In contrast, metabolism- or adipocyte-related pathways were not detected in Cluster A3. Taken together, the findings of the TF motif and pathway analyses indicate a clear correlation of adipocyte-specific gene expression programs with H3K18la, but not with H3K27ac.

### Promoters and enhancers are covered by unique combinations of H3K18la and H3K27ac

Given the distinct distribution of H3K18la- and H3K27ac-marked genomic regions (Figure 1B), we characterized these histone modifications in more detail by separately analyzing promoters and enhancers. To focus on promoter regions, we utilized H3K4me3 data and applied the same clustering method used for H3K18la and H3K27ac (Figure S1A). Here, the H3K4me3-High regions were defined as active promoters and clustered into four groups based on H3K18la and H3K27ac levels (Figure 1E). The proportion of H3K4me3-High promoter regions in Clusters A1–A3 was highly consistent with their distribution on the genome: 32% (3,181/9,812), 20% (346/1,712), and 72% (4,105/5,731) for Clusters A1, A2, and A3, respectively. H3K4me3-High regions were also found in a small population of Cluster A4 (5%, 3,070/56,677). We examined H3K4me3 and ATAC-seq signals in each group (Figures 1E and 1F) and found that these levels were not quantitatively proportional to those of H3K18la and H3K27ac at promoters.

A similar analysis was performed for enhancers, defined by H3K4me3-Low/H3K4me1-High regions (Figures S1A and S1B). We found that 42% of potential enhancer regions were H3K27ac-High (Clusters A1 and A3, 7,806/18,707), 82% of which were co-occupied by H3K18la (6,377/7,806). Importantly, the genomic regions only marked by H3K27ac (Cluster A3) were located more frequently at promoters (72%, see above) than at enhancers (25%, 1,429/5,731). Furthermore, among all putative enhancer regions, only a small fraction was uniquely marked by H3K27ac (Figure S1B, ∼8%, 1,429/18,707). These observations suggest that H3K27ac profiles alone are insufficient to selectively identify the distribution of enhancers.

### A specific set of epigenetic modifiers and TFs occupy H3K18la-marked regions in mESCs

Our analysis revealed that H3K18la and H3K27ac did not essentially co-localize on the genome of mature adipocytes. We wanted to investigate whether H3K18la and H3K27ac exhibit unique distributions in other cell types as well. Hence, we next focused on mESCs, for which a variety of public omics datasets are available. Using H3K18la and H3K27ac CUT&Tag datasets in mESCs^15^, all ATAC-seq peaks^27^ (∼89k regions) were classified into four groups (Figures 2A and S2A, Clusters B1–B4). We found that, similar to mature adipocytes, mESCs exhibited both shared and unique localization of H3K18la and H3K27ac, with some differences between the two cell types. In mESCs, a greater proportion of the peaks were marked exclusively by H3K18la (Cluster B2, 7,136 regions). In addition, H3K4me3 signals were more detected in regions having both H3K18la and H3K27ac (Cluster B1, 11,197 regions), as reflected by their locations on the genome (Figure 2B). Clusters uniquely marked by either H3K18la (B2) or H3K27ac (B3, 5,266 regions) showed preferences in their distributions toward intergenic/intronic and promoter regions, respectively (Figure 2B), and this trend was more marked than that in adipocytes (A2 and A3 in Figure 1B).

**Figure 2.**
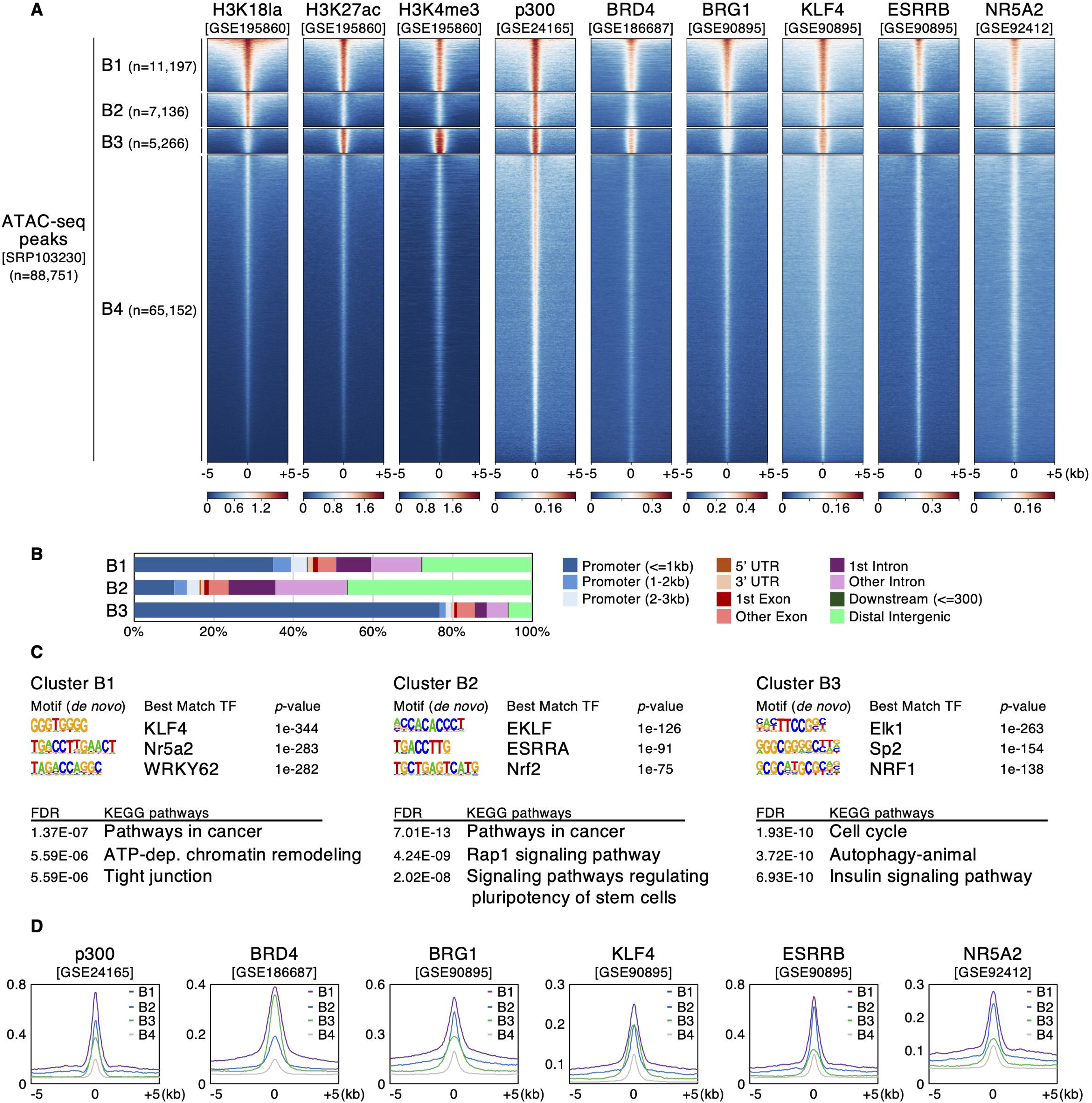
Unique binding profiles of trans-acting factors underlie H3K18la-occupied regions in mESCs. (A) Heatmaps for CUT&Tag and ChIP-seq signals of the indicated modifications and factors at ATAC-seq peak clusters based on H3K18la and H3K27ac levels in mESCs. (B) Genomic distributions of peaks in Clusters B1–B3. (C) TF motif (upper) and KEGG pathway analyses (lower panels) for Clusters B1–B3 (top 3 hits for each analysis). (D) Aggregation plots for binding levels of the indicated factors at each cluster.

TF motif analysis indicated that the Krüppel-like factor (KLF) family– and ESRRB/NR5A2-related sequences were enriched in Clusters B1 and B2 (Figure 2C, upper panels, Table S2). KLF4 is one of the Yamanaka factors key to establishing and maintaining pluripotency^28^, and the orphan nuclear receptors ESRRB and NR5A2 share their recognition sequences and exert cooperative actions on the gene expression network in mESCs^29^. In KEGG pathway analysis using annotated genes (Figure 2C, lower panels, Table S3), Clusters B1 and B2 showed the same pathway as the top hit, and the latter had “Signaling pathways regulating pluripotency of stem cells” in the hit list. These features, which were common between Clusters B1 and B2, were not observed in Cluster B3. Therefore, it is suggested that, similar to that in mature adipocytes, H3K18la marks the genomic regions that represent cell type–specific signatures in mESCs more preferentially than H3K27ac. ChIP-seq data for histone H3 acetylated at lysine 18 (H3K18ac) from two independent studies^7,30^ revealed the distributions distinct from that of H3K18la (Figure S2B), indicating that acetylation and lactylation are differently regulated on the same lysine residue. H3K18la profiles were largely maintained in two different culture conditions/cell states (Figure S2C, Primed and Naïve), supporting the idea that this histone mark is tightly linked to the nature intrinsic to cell identity.

We then explored ChIP-seq data for epigenetic modifiers and TFs identified in the motif analysis^3,28,31,32^ to examine their associations with H3K18la and H3K27ac (Figures 2A and 2D). p300 is a major histone acetyltransferase that also catalyzes histone lactylation^14^. Consistently, p300 localized best at the regions covered by both H3K18la and H3K27ac (Cluster B1), and its binding was also observed, to a lesser degree, in the individually marked regions (B2 and B3). The distribution of BRD4, a transcriptional coactivator that binds to acetylated histones through its bromodomains^33^, was correlated with that of H3K27ac (Clusters B1 and B3). In contrast, BRG1, the ATPase subunit of the SWI/SNF chromatin remodeling complex, was preferentially found in the H3K18la-marked regions (Clusters B1 and B2), which might reflect its potential role as a lactylated histone reader^34^. Among the TFs identified in the motif analysis, ESRRB and NR5A2 exhibited good correlation with H3K18la in their localization, as expected, whereas KLF4 binding was found in Cluster B3 in addition to B1 and B2. We speculate that this could be a result of the motif-independent binding of KLF4 assisted by other TFs^28^. Indeed, ChIP-seq profiles of OCT4 and SOX2, which co-occupy the binding sites with KLF4^28^, were analogous to that of KLF4 (Figures S2D and S2E). NANOG was more found at H3K18la-marked regions like ESRRB and NR5A2, while the binding of MYC, which is pivotal but not specific to mESCs, was associated with H3K27ac (Figures S2D and S2E). Overall, our analysis of mESCs, together with that of adipocytes, highlights the unique cell type–specific roles of H3K18la.

### H3K18la identifies p300/CBP-dependent cis-regulatory elements even without H3K27ac co-occupancy

Using A-485, a potent selective inhibitor of p300/CBP, Narita et al. generated a large series of omics datasets and reported how epigenetic and transcriptional landscapes are dynamically shaped by the activity of these enzymes in mESCs^6^. We used this valuable data to investigate the p300/CBP-mediated regulation of H3K18la- and H3K27ac-occupied regions. We first focused on H3K4me3-High active promoter regions that were clustered based on H3K18la and H3K27ac levels (Figures 3A and S2A). ATAC-seq signals both in the presence and absence of A-485 treatment were analyzed in these clusters, and the effect of the treatment was evaluated using the log2 fold change (log2FC) between the two conditions (Figures 3A and 3B). The promoters in both Clusters B1 and B3 had the highest chromatin accessibilities; however, as is evident from the log2FC values, they exhibited differential sensitivities against A-485. The reduction in ATAC-seq signals upon p300/CBP inhibition was most significant in Cluster B1, whereas the responsiveness of Cluster B3 was marginal and comparable to that of Cluster B4, where both H3K18la and H3K27ac marks were absent. It should be noted that the promoters at Cluster B2, despite being minor, indicated a greater sensitivity to A-485 than those at Cluster B3. To examine the effect of A-485 treatment on transcription from promoters, we analyzed changes in RNA polymerase II (Pol II) and nascent transcript levels on the gene bodies of each cluster using ChIP-seq and 5-ethynyluridine sequencing (EU-seq) data (Figures 3C and 3D). In both datasets, Clusters B1 and B2 showed reduced transcriptional activity upon p300/CBP inhibition, similar to decreased accessibility at the promoter regions. At Cluster B4, on average, hardly any effect was observed on ongoing transcription. Surprisingly, A-485 treatment increased Pol II and nascent transcript levels of Cluster B3 genes, even though their promoters were marked by H3K27ac.

**Figure 3.**
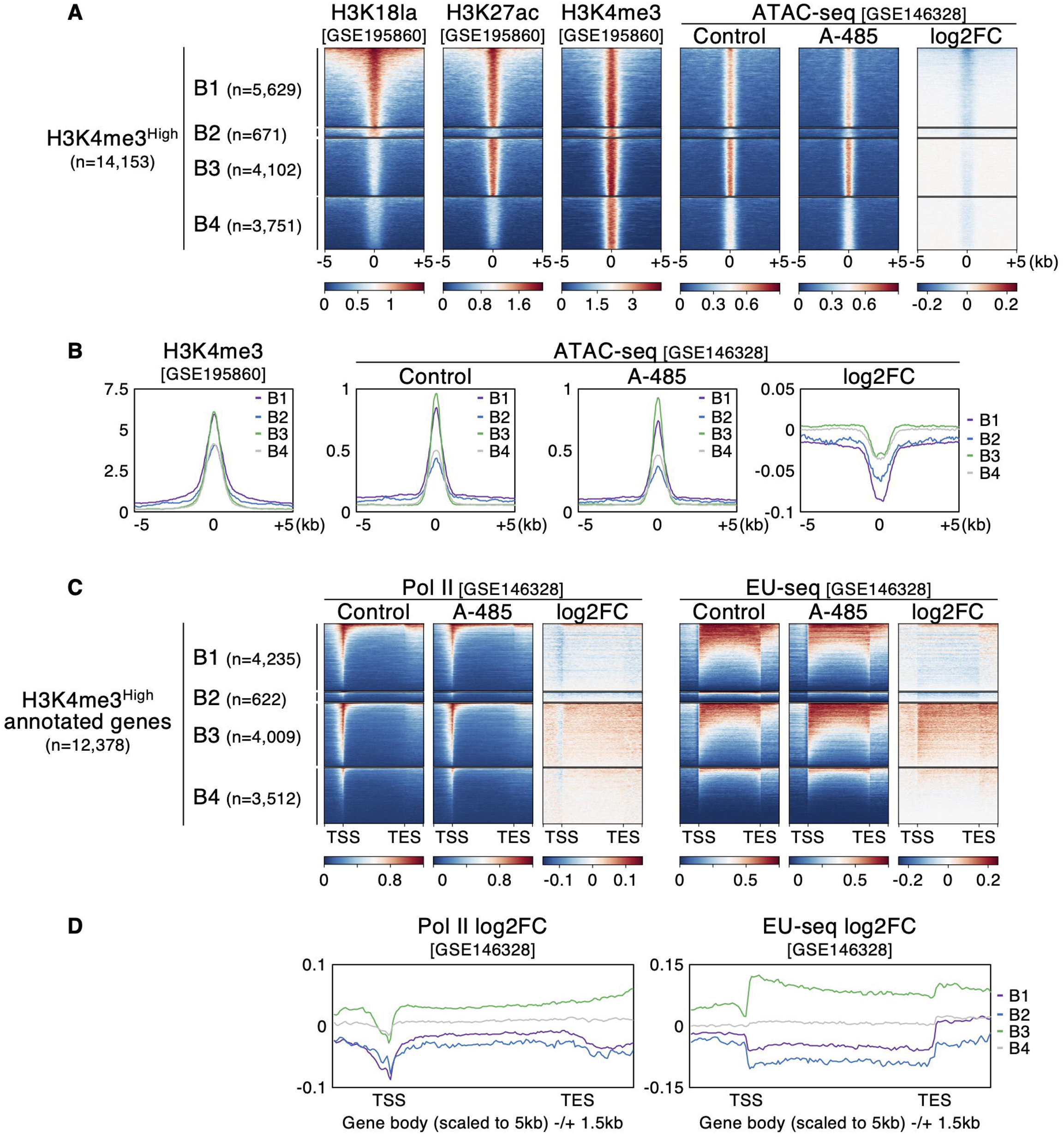
H3K27ac-marked promoters are p300/CBP-dependent only when co-occupied with H3K18la. (A) Heatmaps for CUT&Tag and ATAC-seq signals at H3K4me3-High regions of each cluster in mESCs. Effects of A-485 treatment on ATAC-seq levels are visualized using log2FC values (Control versus A-485). (B) Aggregation plots for the indicated data at H3K4me3-High regions of each cluster. (C) Heatmap presentation of Pol II loading (ChIP-seq) and nascent transcript levels (EU-seq) at gene bodies annotated with H3K4me3-High regions in each cluster. Signals for gene bodies (scaled to 5kb) -/+ 1.5kb are shown. (D) Aggregation plots for log2FC values of Pol II ChIP-seq and EU-seq at gene bodies of each cluster.

The effects of p300/CBP inhibition on enhancers were also analyzed using the same strategy; however, the focus was on H3K4me3-Low/H3K4me1-High regions, as defined using their CUT&Tag and ChIP-seq data^15,35^ (Figures 4 and S2A). Similar to what was observed with the promoter regions, we found a reduction in chromatin openness (Figures 4A and 4B) and transcriptional suppression of annotated genes (Figures 4C and 4D) at Clusters B1 and B2 upon p300/CBP inhibition. In contrast, A-485 treatment did not show any inhibitory (or activating) effects on Cluster B3 enhancers. To gain further insight into the functionalities of H3K18la- and/or H3K27ac-marked enhancers, we integrated public Self-Transcribing Active Regulatory Region sequencing (STARR-seq) data of mESCs^36^, which enables the unbiased identification of cis-regulatory elements and measurement of their enhancer potentials through massively parallel reporter assays. The enhancers at Clusters B1 and B2 uniformly had high STARR-seq signals; however, the levels at Cluster B3 were negligible (Figures 4A and 4B), indicating a clear correlation between H3K18la and enhancer activity. Collectively, these findings suggest that the genomic regions occupied by H3K18la serve as functional active enhancers independent of H3K27ac, although these modifications can co-localize at certain promoter regions (Cluster B1 in Figure 3). These results also indicate that p300/CBP-mediated epigenomic regulations have a stronger association with H3K18la. Intriguingly, when analyzing H3K27ac levels in the presence or absence of A-485 treatment, we observed differential sensitivities to p300/CBP inhibition at Clusters B1 and B3 (Figures S2F and S2G), suggesting locus-specific regulation on this histone mark.

**Figure 4.**
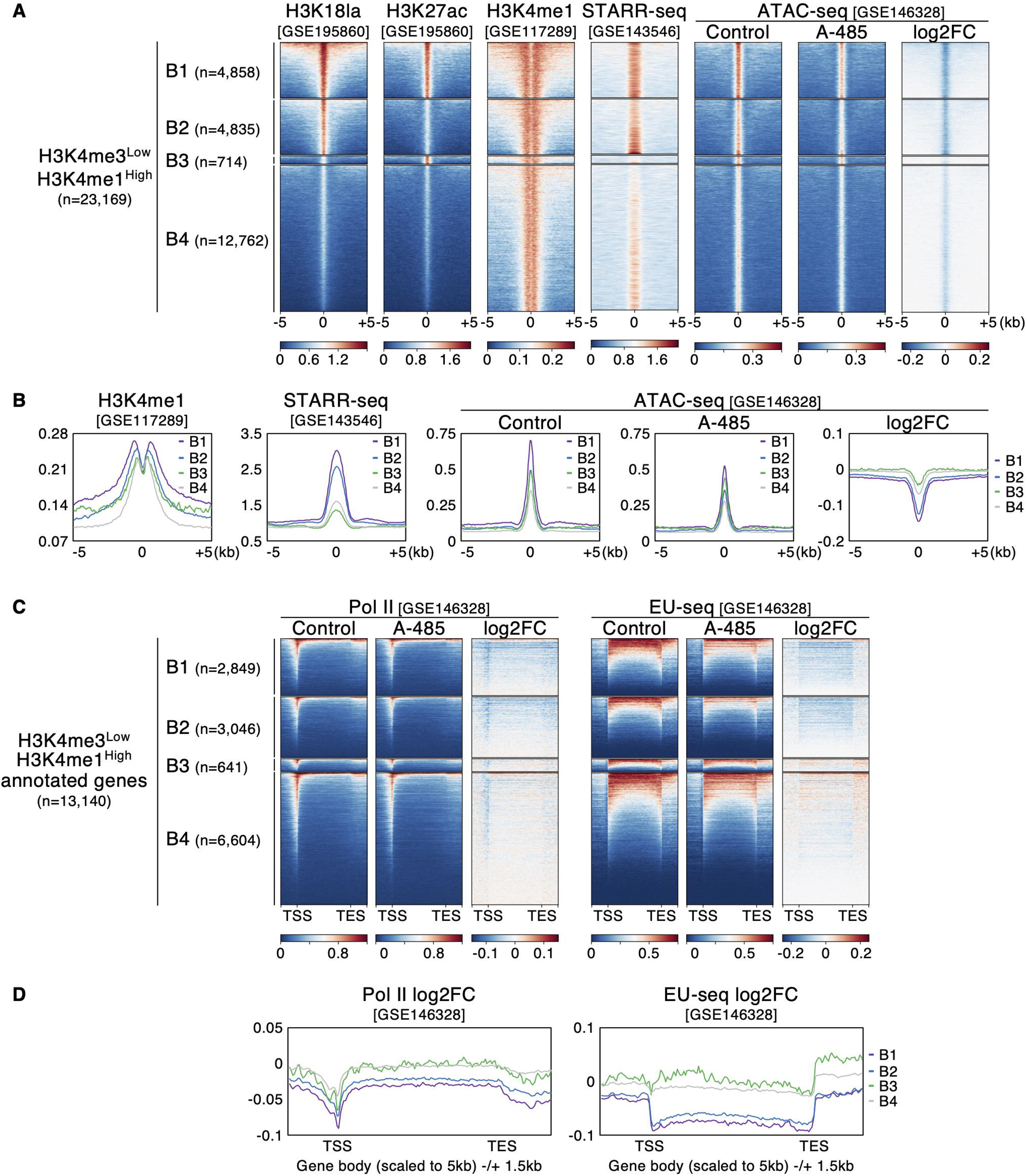
H3K18la-bound enhancers are functional and actively regulated by p300/CBP, irrespective of H3K27ac presence. (A) Heatmaps of CUT&Tag, ChIP-seq, STARR-seq, and ATAC-seq signals at H3K4me3-Low/H3K4me1-High regions of each cluster in mESCs. (B) Aggregation plots for the indicated data at H3K4me3-Low/H3K4me1-High regions of each cluster. (C) Heatmap presentation of Pol II ChIP-seq and EU-seq signals at gene bodies annotated with H3K4me3-Low/H3K4me1-High regions in each cluster. (D) Aggregation plots for log2FC values of Pol II ChIP-seq and EU-seq at gene bodies of each cluster.

### H3K27ac-occupied regions are associated with transcriptional regulations that involve liquid droplet formation

Super-enhancers are thought to be formed by the LLPS of BRD4 and MED1, by which multiple enhancers assemble at nuclear condensates^11^. Although Clusters B1 and B2 were associated with enhancer activity, only the former was bound by BRD4 (Figure 2A and 2D). Thus, where other factors involved in super-enhancers were located on the genome was investigated. To determine this, ChIP-seq profiles of the cohesin subunit SMC1^37^ and the Mediator subunit MED1^38^ were analyzed (Figure 5A and 5B). SMC1 was distributed throughout all clusters, whereas MED1 indicated selective binding to Clusters B1 and B3, similar to BRD4. Based on these results, we hypothesized that, although functional (typical) enhancers were enriched in Clusters B1 and B2, super-enhancers were associated more specifically with Cluster B1. To test this possibility, super-enhancer regions were determined using MED1 ChIP-seq data (1,062 regions), and their overlap with Clusters B1–B4 was examined (Figure 5C). As expected, Cluster B1 had the highest overlap ratio with the super-enhancers. Consistently, the super-enhancers located at *Klf4*, *Pou5f1* (encoding OCT4), and *Nanog* loci were covered by both H3K18la and H3K27ac (Figure 5D). In contrast, Cluster B3 showed limited overlap which was comparable to that of Cluster B2 (Figure 5C), suggesting that the binding of BRD4 and MED1 is important but not sufficient for super-enhancer formation.

**Figure 5.**
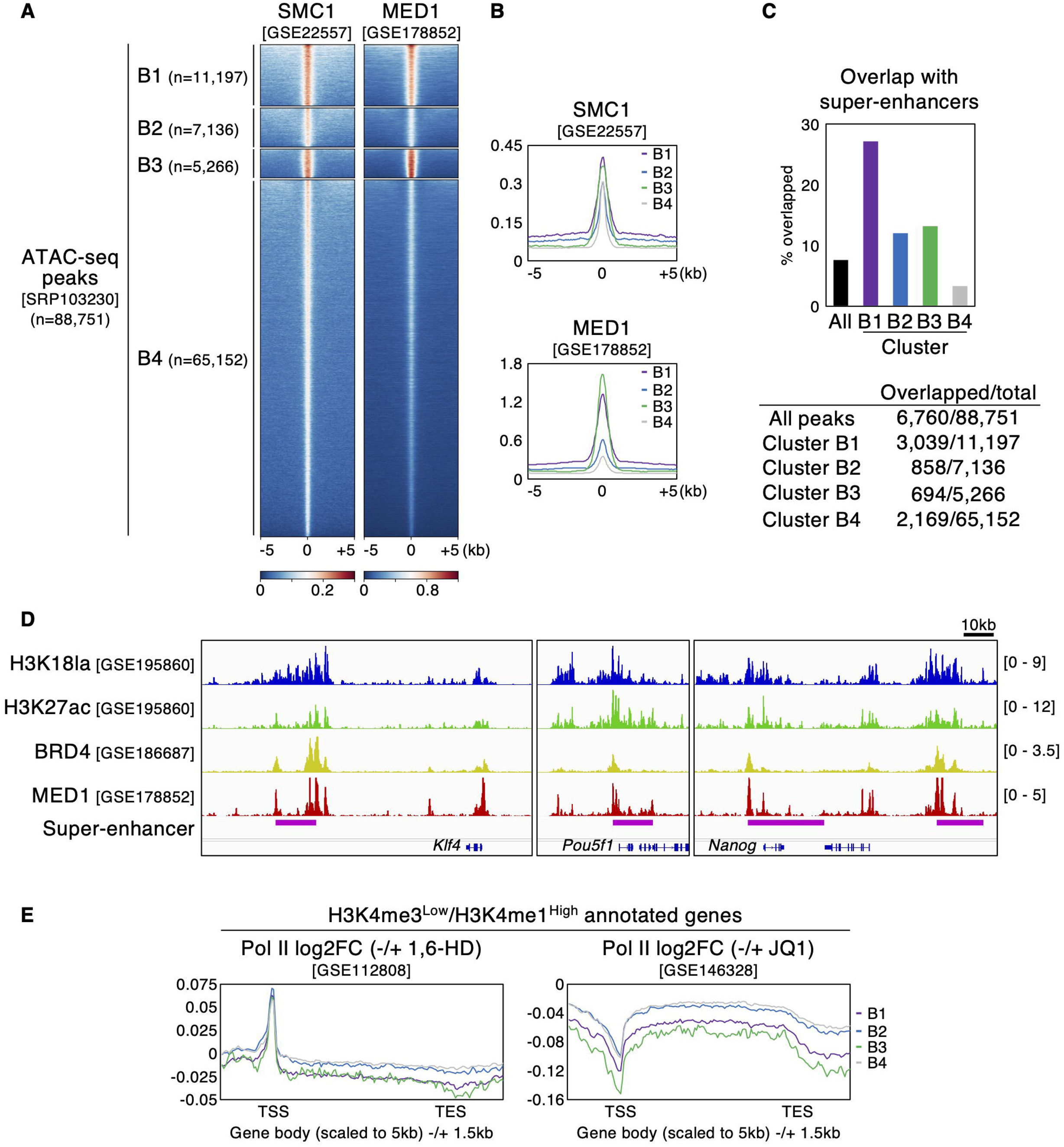
H3K18la-High/H3K27ac-High regions are enriched in super-enhancers. (A-B) Heatmaps (A) and aggregation plots (B) of ChIP-seq signals of SMC1 and MED1 at Clusters B1–B4. (C) Overlaps between each cluster and super-enhancers. Upper and lower panels indicate % overlapped and peak numbers, respectively. (D) Genome browser tracks of the indicated data at super-enhancers close to *Klf4*, *Pou5f1*, and *Nanog* genes. (E) Aggregation plots for log2FC values of Pol II ChIP-seq with or without 1,6-HD (left) or JQ1 treatments (right) at gene bodies annotated with H3K4me3-Low/H3K4me1-High regions in each cluster.

Liquid droplets of BRD4 and MED1 are dissolved upon 1,6-hexanediol (1,6-HD) treatment, which disrupts hydrophobic interactions^11^. Similarly, JQ1, a small-molecule inhibitor of bromodomain and extra-terminal proteins including BRD4, suppresses the formation of BRD4 condensates^39^. BRD4 inhibition reduces MED1 binding, especially to super-enhancers^40,41^ (Figure S3A and S3B). Hence, Pol II ChIP-seq datasets with or without 1,6-HD^11^ or JQ1 treatments^6^ were analyzed to investigate the effects of condensate disruption on enhancer-driven transcription (Figure 5E). Similar to the analysis of p300/CBP inhibition (Figure 4D), Pol II levels at the gene bodies annotated with putative enhancers in Clusters B1–B4 were examined, and the effects of the treatments were visualized using log2FC values. Although 1,6-HD and JQ1 affected Pol II occupancy at transcription start sites (TSSs) oppositely, both treatments decreased the elongating Pol II levels at the gene bodies in Clusters B1 and B3, where H3K27ac was present. In contrast, Clusters B2 and B4 showed limited responses to the treatments, indicating that although H3K18la-marked regions exhibited intrinsic enhancer activity, H3K18la occupancy alone was insufficient to support gene expression involving LLPS. Taken together, these results suggest that the H3K27ac-covered regions are responsible for the cooperative regulation of multipartite enhancers mediated by LLPS. This relationship between H3K27ac and LLPS-driven enhancer cooperativity is similar to that between H3K18la and p300/CBP-dependent enhancer activity, indicating differential functions of these modifications.

### H3K18la and H2BNTac share unique features as active enhancer marks but exhibit distinct genomic distributions

The characteristics of H3K18la, such as its close association with enhancers and its p300/CBP-dependent regulation, are highly analogous to those of H2BNTac, a recently reported signature of active enhancers^4^. Therefore, we compared the genomic distributions of H3K18la and H2BNTac (Figure 6). Using ChIP-seq data for multiple acetylated lysine residues in H2B (K5, K12, K16, and K20), we examined their levels at the four clusters based on H3K18la and H3K27ac (Figure 6A). We observed a certain similarity between the localization of H3K18la and that of H2BNTac (e.g., their absence at Cluster B3); however, we also observed some differences (e.g., H2BNTac signals at Cluster B4). To directly verify the common and distinct genomic distributions of H3K18la and H2BNTac, ATAC-seq peaks were clustered into four groups, as was done for H3K18la and H3K27ac, using H2BK16ac data as a representative (Figures S4A and S4B). As expected, we identified a significant number of regions with single and dual H3K18la and H2BK16ac marks. The same clustering scheme was then adopted using not only H3K18la and H2BK16ac, but also H3K27ac CUT&Tag/ChIP-seq data, resulting in eight groups: Clusters C1 and C2 for H3K18la-High/H2BK16ac-High, C3 and C4 for H3K18la-High/H2BK16ac-Low, C5 and C6 for H3K18la-Low/H2BK16ac-High, C7 and C8 for H3K18la-Low/H2BK16ac-Low—odd numbers for H3K27ac-High, and even numbers for H3K27ac-Low (Figure 6B). Except for Cluster C5 (H3K18la-Low/H2BK16ac-High/H3K27ac-High, 450 regions), several thousands of the regions were found at each cluster having at least one modification (2.3k-8.4k regions), confirming distinct distributions and combinations of the individual marks. We analyzed the responsiveness of each cluster to p300/CBP inhibition using the ATAC-seq data, with or without A-485 treatment (Figures 6C and 6D). Consistent with an earlier analysis, the sensitivities to A-485 were largely independent of H3K27ac occupancy (e.g., Clusters C1 versus C2). Clusters C1–C2 were the most sensitive to A-485, whereas C3–C4 and C5–C6 exhibited modest and comparable reductions in chromatin accessibility upon p300/CBP inhibition. This suggests that H3K18la and H2BNTac may play an additive or synergistic role in determining p300/CBP dependency at their marked regions.

**Figure 6.**
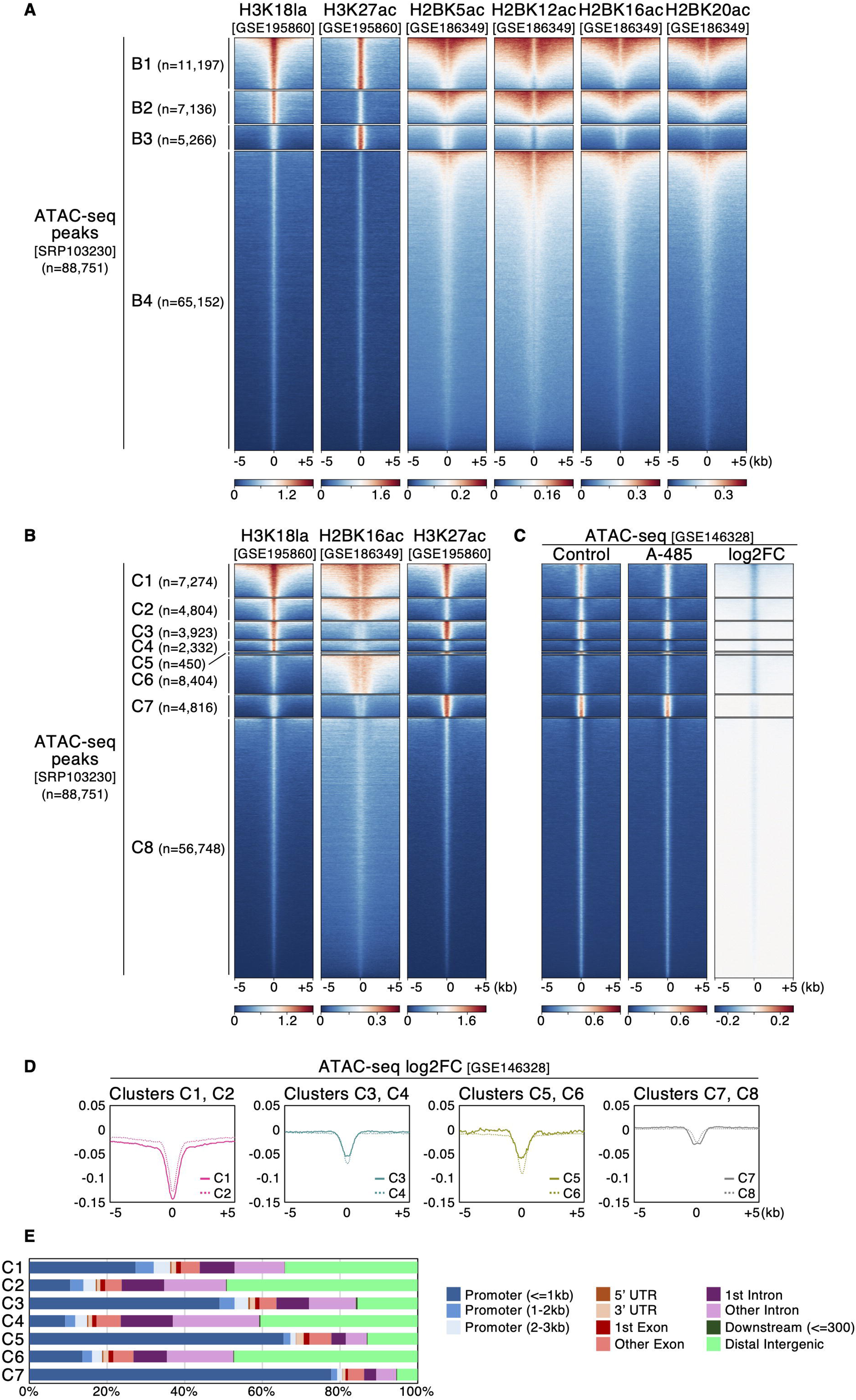
H3K18la and H2BNTac occupy p300/CBP-regulated regions both uniquely and in combination. (A) Heatmaps of CUT&Tag and ChIP-seq signals of the indicated modifications at Clusters B1–B4 in mESCs. Note that genomic regions are shown in descending order of H2BNTac levels. (B) Heatmap visualization of H3K18la, H2BK16ac, and H3K27ac signals at Clusters C1–C8, classified based on those levels (descending order based on all three modifications). (C) Heatmaps of ATAC-seq signals with or without A-485 treatment and log2FC values at Clusters C1–C8 (the same order as in B). (D) Aggregation plots of ATAC-seq log2FC values at Clusters C1–C8. (E) Distributions of Clusters C1–C7 in the genome.

Finally, the genomic distribution of each cluster was analyzed (Figure 6E), revealing three general trends. First, the genomic loci where H3K18la, H2BK16ac, and H3K27ac all coexisted (Cluster C1) were located at both promoters (∼40%) and intergenic/intronic regions (∼60%). Second, the regions marked by H3K18la and/or H2BK16ac, but not H3K27ac (Clusters C2, C4, and C6), had limited localization at promoters (less than 20%). Third, H3K27ac-occupied regions were conversely found to be much more prevalent at promoters (50–80%, Clusters C3, C5, and C7). Altogether, our analysis strongly indicates that H3K18la, along with H2BNTac, serves as the most effective marker for functional enhancers actively regulated by p300/CBP, with shared and unique distributions in the genome.

## DISCUSSION

Using a variety of public multi-omics datasets, we systematically analyzed H3K18la distributions in the genome and uncovered unique features that differed from those of H3K27ac. In both mature adipocytes and mESCs, H3K18la and H3K27ac were found to be mainly localized at intergenic/intronic and promoter regions, respectively, and only the former was associated with cell type–specific TF binding. Although it is well accepted that H3K27ac occupies active enhancers^3^, only a limited number of enhancers were uniquely occupied by H3K27ac in mESCs with limited enhancer activity, as indicated by STARR-seq data. Instead, H3K18la was found to be a more reliable marker of functional enhancers. Of all the putative enhancers, more than 40% of the regions were H3K18la-High, with a robust enhancer potential and sharp p300/CBP dependency, regardless of H3K27ac co-occupancy. Thus, the widely adopted H3K27ac-based analysis does not efficiently identify the genome-wide distribution of active enhancers, suggesting a need to reconsider the current framework of epigenome studies.

The presence of H3K27ac was not limited to promoters; its localization was detected in a subset of enhancers marked by H3K18la. The distribution of BRD4 and MED1, as well as their ability to regulate transcription by forming liquid droplets, were highly correlated with H3K27ac localization. Thus, it is suggested that while H3K18la is a marker of whether a given locus acts as an enhancer, H3K27ac occupies the genomic regions responsible for how multiple cis-regulatory elements behave cooperatively through LLPS. The observation that super-enhancers best overlapped with genomic regions covered by both H3K18la and H3K27ac supports this notion. Based on these findings, we propose a model in which “enhancer potential” associated with H3K18la and “enhancer cooperativity” linked to H3K27ac are two independent, separable properties, and when combined, achieve synergistic enhancer activation (i.e., super-enhancers). This model is in line with the study on the alpha-globin super-enhancer showing that not only typical enhancers, but also facilitators, are necessary to fully activate gene expression^13^. Recently, Narita et al. reported that H2BNTac occupies the vast majority of active enhancers in mESCs and that H2BNTac intensity best predicts p300/CBP-target genes^4^. Our analysis revealed notable similarities between H3K18la and H2BNTac. First, as mentioned above, H3K18la distribution was highly biased toward intergenic/intronic regions, where enhancers predominantly reside. Second, we observed that transcriptional outputs from H3K18la-marked promoters and enhancers were sensitive to the p300/CBP inhibitor, A-485. Third, the regions covered by either H3K18la or H2BK16ac revealed a comparable reduction in chromatin accessibility upon p300/CBP inhibition. The presence of singly marked regions also indicates that H3K18la and H2BNTac can be distributed and regulated individually in the genome. Independent studies have revealed that metabolic remodeling upon changes in glucose or lactate levels has a greater impact on histone lactylation than on acetylation^14,42^. Therefore, we speculate that H3K18la has shared features with H2BNTac (such as enhancer marking), as well as unique ones, which mediate metabolism-driven gene expression.

Although our study and the work by the Choudhary group^4^ establish H3K18la and H2BNTac as the so-far best markers for active enhancers, these modifications were also found in some promoters co-occupied by H3K27ac. This suggests that it is too simple to characterize genomic regions of interest using a single or a couple of histone marks. Given the virtually unlimited combinations of histone modifications that have been identified to date, further studies using large datasets of histone marks are needed to identify more functional histone codes that define individual genomic elements such as enhancers. It is also important to discriminate the functions of histone modifications *per se* from those of their marked genomic regions. Although we have elucidated the unique features of H3K18la- and H3K27ac-marked regions, our findings do not exclude the possibility that these modifications themselves are dispensable for enhancer regulation, as reported previously^7^.

One of the major issues that remain to be addressed is identifying the enzyme(s) that catalyzes H3K18la at each locus. Several reports have demonstrated p300-mediated histone lactylation^14,43,44^, which is consistent with our results showing the exclusive dependency on this enzyme at H3K18la-marked regions. Other studies identified KAT2A/GCN5^45^ and KAT7/HBO1^46^ as histone lactyltransferases. Furthermore, recent studies have revealed non-canonical functions of class I histone deacetylases HDAC1-3 that catalyze histone lactylation^42,47^, although they have also been shown to act as histone delactylases^48^. We examined HBO1 ChIP-seq data in mESCs^49^ and found that its localization was more strongly associated with H3K27ac than with H3K18la (Figures S5A and S5B). ChIP-seq datasets for HDACs^50,51^ indicated that HDAC3, rather than HDAC1 and 2, correlated with H3K18la in their distribution (Figures S5C and S5D). H3K27ac levels were largely unaffected by *Hdac3* knockout in mESCs^51^ (Figures S5E and S5F), suggesting an H3K27ac-independent role for HDAC3. Interestingly, a study on *Drosophila* revealed that depletion of Nej/p300 and HDAC3 reduces histone lactylation in oocytes^52^. Concordantly, another study revealed that the inhibition of not only p300/CBP but also class I HDACs resulted in a decrease in histone lactylation levels, including H3K18la^44^. These lines of evidence imply that p300 and/or HDAC3 may be involved in H3K18la deposition in the mESC genome, although this needs to be carefully addressed in future studies. In addition, the fact that a single enzyme (e.g., p300) can catalyze both histone lactylation and acetylation presents further challenges in advancing our understanding of the roles of individual modifications. Thus, the development of novel experimental tools, such as selective inhibitors of each histone mark, is highly desired.

In summary, this study identified H3K18la as a histone mark for p300/CBP-regulated cis-regulatory elements. Our analysis also revealed that H3K27ac-marked regions are responsible for enhancer cooperativity, rather than potential, that involves LLPS. We believe that our findings regarding the significant differences between H3K18la and H3K27ac will greatly help update the current understanding of how enhancer landscapes are shaped by histone modifications.

## METHODS

### Data processing

All datasets used in this study were obtained from public databases, as summarized in Table S1. Except for STARR-seq (see below), sequencing data were processed from the original FASTQ files on Galaxy (https://usegalaxy.org/ and https://usegalaxy.eu/)^53^ as follows: Using Download and Extract Reads in FASTQ (Galaxy Version 3.1.1+ galaxy1), FASTQ files were downloaded from the Sequence Read Archive of the National Center for Biotechnology Information (NCBI SRA). After preprocessing with fastp (Galaxy Version 1.1.0+ galaxy0), the reads were mapped to the mouse mm9 reference genome using Bowtie2 (Galaxy Version 2.5.4+ galaxy0, --very-sensitive). Mitochondrial DNA reads were removed using Filter SAM or BAM, output SAM or BAM (Galaxy Version 1.8+ galaxy1). Multi-mapped and duplicate reads were omitted using Samtools sort (Galaxy Version 2.0.8), fixmate (Galaxy Version 1.22+ galaxy1, -m), view (Galaxy Version 1.22+ galaxy1, -q 40 -F 0×4), and markdup (Galaxy Version 1.22+ galaxy1, -r). The resulting BAM files were converted to bigWig files using the bamCoverage function of deepTools (Galaxy Version 3.5.4+ galaxy0, --normalizeUsing CPM). Where applicable, replicate data for each condition were merged using deepTools bigwigAverage (Galaxy Version 3.5.4+ galaxy0). The bigwigCompare function of deepTools (Galaxy Version 3.5.4+ galaxy0) was used to calculate log2FC values. Except for Figure S2C, the analysis on H3K18la in mESCs was performed using CUT&Tag data for the Primed (Serum LIF) condition. Visualization of bigWig data was done on Integrative Genomics Viewer (IGV, version 2.19.1).

The author-processed STARR-seq data^36^ (serum+LIF condition, bigWig files for the Empirical Bayes shrunken enrichment) were retrieved from the NCBI Gene Expression Omnibus (GEO, GSE143546), and the replicates were merged as described above for subsequent analysis.

### Peak-calling for ATAC-seq data

Processed BAM files of ATAC-seq data were subjected to MACS2 callpeak (Galaxy Version 2.2.9.1+ galaxy0) with the following options: --format BAMPE --gsize mm --nomodel –qvalue 0.05 --broad --broad-cutoff 0.05 --keep-dup all. Output bed files from replicates were concatenated (cat), sorted with bedtools SortBED (Galaxy Version 2.31.1+ galaxy0), and merged using bedtools MergeBED (Galaxy Version 2.31.1+ galaxy2). The resulting bed files were used as master peak lists.

### Generation of heatmaps and aggregation plots

To perform k-means clustering of ATAC-seq peaks based on each histone modification (k=2, High and Low groups), ChIP-seq and CUT&Tag signals around all ATAC-seq peaks were analyzed using the deepTools computeMatrix (Galaxy Version 3.5.4+ galaxy0, reference-point --referencePoint center --beforeRegionStartLength 1500 --afterRegionStartLength 1500 --missingDataAsZero) and plotHeatmap (Galaxy Version 3.5.4+ galaxy0, --outFileSortedRegions --kmeans 2). Using outputs from plotHeatmap, bed files for High and Low groups were generated. To combine the High and Low groups for multiple histone modifications, bed files were subjected to bedtools Intersect intervals (Galaxy Version 2.31.1+ galaxy0, -wa), SortBED, and MergeBED. Heatmap visualization of omics datasets at each group/cluster was carried out using the deepTools computeMatrix (reference-point --referencePoint center --beforeRegionStartLength 5000 --afterRegionStartLength 5000 --missingDataAsZero) and plotHeatmap. Unless otherwise specified, genomic regions in each group were sorted in descending order (--sortRegions descend) based on H3K18la and H3K27ac levels (--sortUsingSamples). Aggregation plots were generated using the plotProfile function of deepTools (Galaxy Version 3.5.4+ galaxy0).

For visualizing Pol II ChIP-seq and EU-seq data at gene bodies, bed files for each gene (based on the longest transcripts), from TSS to transcription end site (TES), were generated as described previously^54^ and used for computeMatrix (scale-regions --regionBodyLength 5000 --beforeRegionStartLength 1500 --afterRegionStartLength 1500 --missingDataAsZero), plotHeatmap (in descending order based on Pol II ChIP-seq and EU-seq signals for control samples), and plotProfile of deepTools.

### Peak annotation and KEGG pathway and TF motif analyses

Peak annotation with nearest genes and examination of genomic locations were performed using ChIPseeker (Galaxy Version 1.28.3+ galaxy0) using a GTF file for the mm9 genome obtained from the University of California, Santa Cruz (UCSC) website (mm9.refGene.gtf, https://hgdownload.soe.ucsc.edu/goldenPath/mm9/bigZips/genes/). KEGG pathway analysis of gene sets was conducted using ShinyGO 0.85.1 (https://bioinformatics.sdstate.edu/go/).

TF motif analysis was performed using the findMotifsGenome.pl of HOMER (version 4.11.1) using ATAC-seq master peak lists as the background (-bg).

### Identification of super-enhancers

ChIP-seq data for MED1 were processed as described above. The resultant BAM files of replicate data were merged using Samtools merge (Galaxy Version 1.22+galaxy2) and converted to a SAM file with Samtools view (version 1.18). The SAM file was subjected to makeTagDirectory and findPeaks (-style super) of HOMER (version 4.11.1) to identify super-enhancer regions. Overlaps with super-enhancers were analyzed using bedtools Intersect intervals.

## ACKNOWLEDGMENTS

We thank Dr. Mitsuru Okuwaki (Kitasato University) and all the members of the Inagaki laboratory for helpful discussions. We would also like to thank Editage (www.editage.jp) for English language editing. This work was supported by JSPS KAKENHI (Grant Numbers 23K07983, 24H01222, and 26K11422 to T.K., and 23K27647 to T.I.), the Gunma Foundation for Medicine and Health Science, the Suzuken Memorial Foundation, the Shimadzu Science Foundation, Kurata Grants by The Hitachi Global Foundation, the Research Foundation for Opto-Science and Technology, and the Takeda Science Foundation to T.K.

## AUTHOR CONTRIBUTIONS

Conceptualization, T.K.; methodology, T.K.; investigation, T.K.; formal analysis, T.K.; data curation, T.K.; visualization, T.K.; funding acquisition, T.K. and T.I.; project administration, T.I.; supervision, T.I.; writing – original draft, T.K.; writing – review & editing, T.I.

## DECLARATION OF INTERESTS

The authors declare no competing interests.

## FIGURE LEGENDS

**Figure S1.**
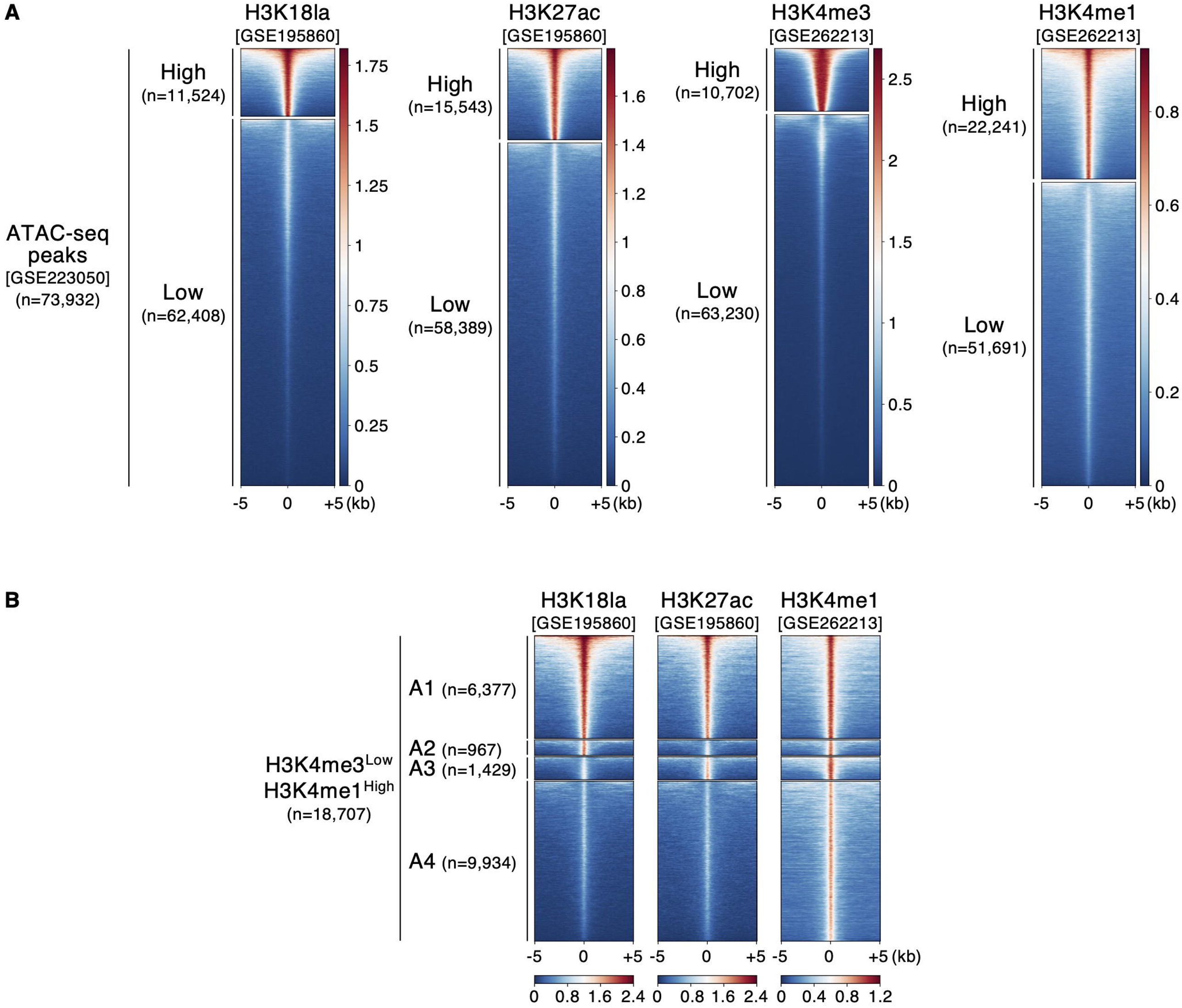
Clustering of adipocyte ATAC-seq peaks based on histone modifications. (A) Heatmaps of CUT&Tag signals of the individual modifications at ATAC-seq peaks after k-means clustering (k=2) in mature adipocytes (peak center -/+ 5kb, descending order in each modification). (E) Heatmap visualization of signals for the indicated data at H3K4me3-Low/H3K4me1-High regions of Clusters A1–A4.

**Figure S2.**
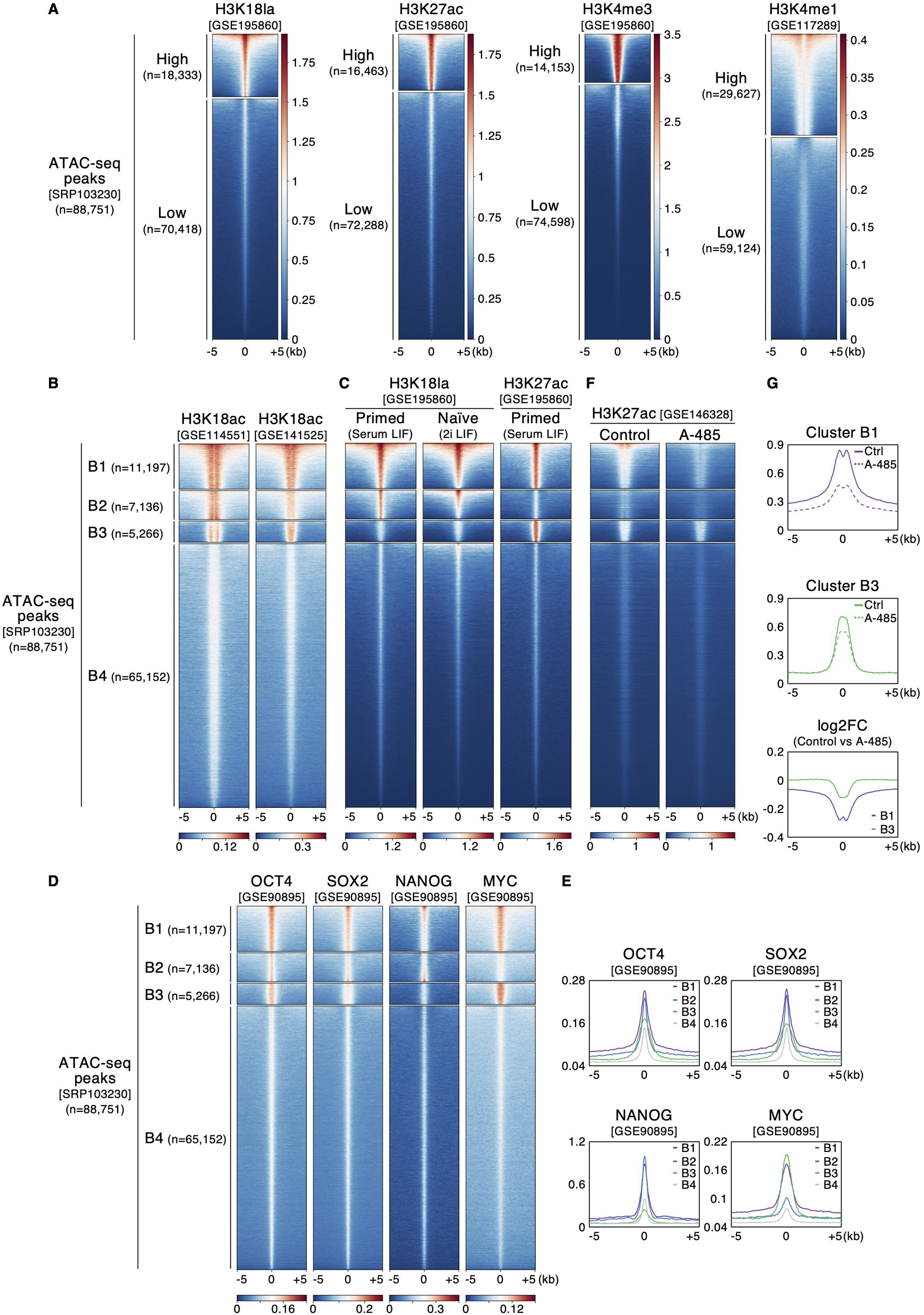
Distribution of histone modifications and their changes under certain culture conditions in mESCs. (A) Heatmaps of CUT&Tag and ChIP-seq signals of the individual modifications at ATAC-seq peaks, after k-means clustering (k=2) in mESCs (descending order in each modification). (B) Heatmap visualization of two independent H3K18ac ChIP-seq data in mESCs. (C) Heatmap visualization of H3K18la and H3K27ac signals in mESCs under the two different culture conditions: Primed (Serum LIF) and Naïve (2i LIF). (D) Heatmaps for ChIP-seq signals of the indicated TFs at each cluster. (E) Aggregation plots for binding levels of the indicated TFs at each cluster. (F) Heatmaps of H3K27ac signals with or without A-485 treatment at Clusters B1–B4. (G) Aggregation plots of H3K27ac levels and log2FC values at Clusters B1 and B3.

**Figure S3.**
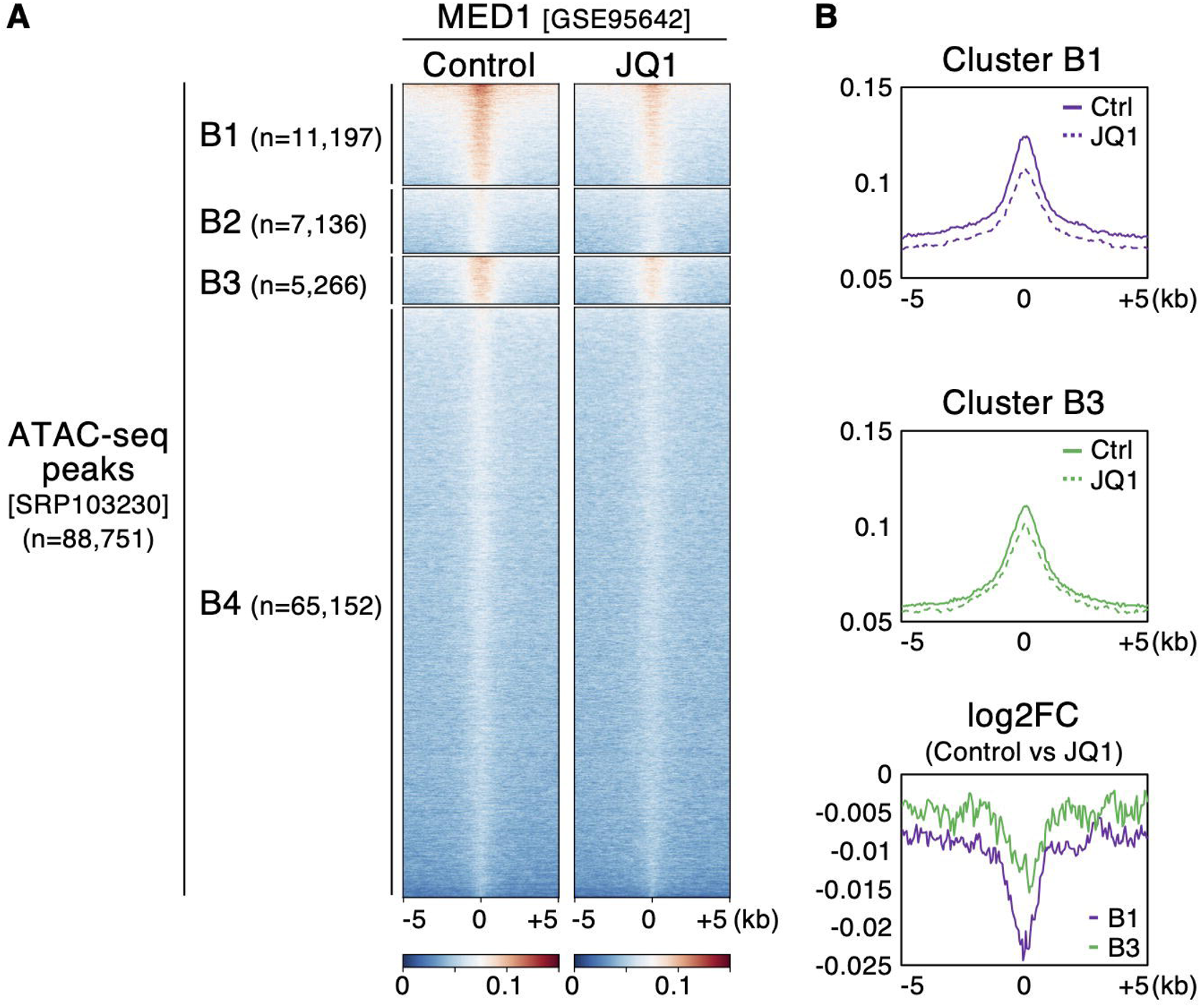
JQ1 treatment attenuates MED1 binding. (A) Heatmaps of MED1 signals with or without JQ1 treatment at Clusters B1–B4. (B) Aggregation plots of MED1 levels and log2FC values at Clusters B1 and B3.

**Figure S4.**
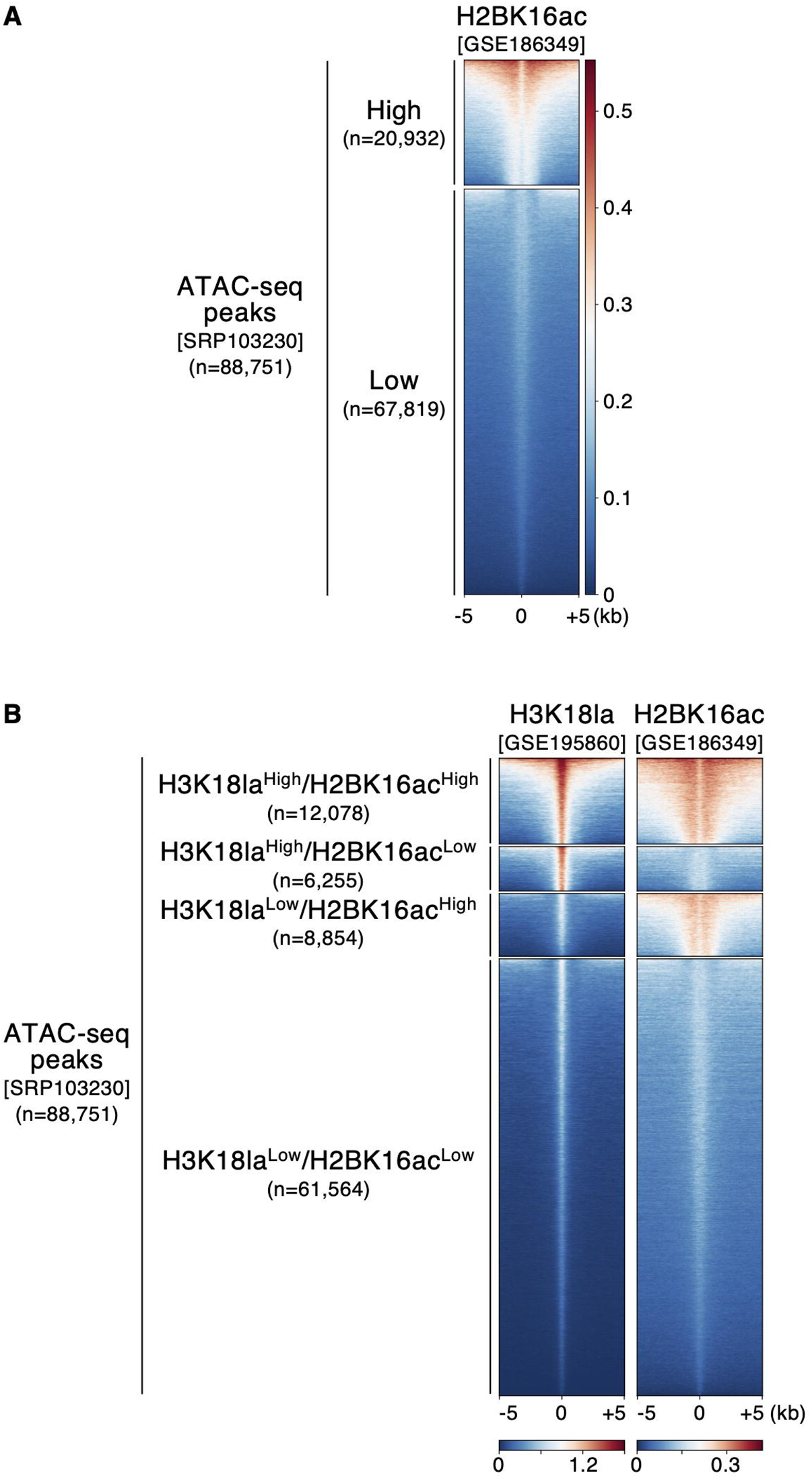
H2BK16ac localization on the genome in mESCs. (A) A heatmap of H2BK16ac ChIP-seq signals at ATAC-seq peaks with k-means clustering (k=2) in mESCs (descending order). (B) Heatmap visualization of H3K18la and H2BK16ac levels at the four clusters based on those levels (descending order based on both modifications).

**Figure S5.**
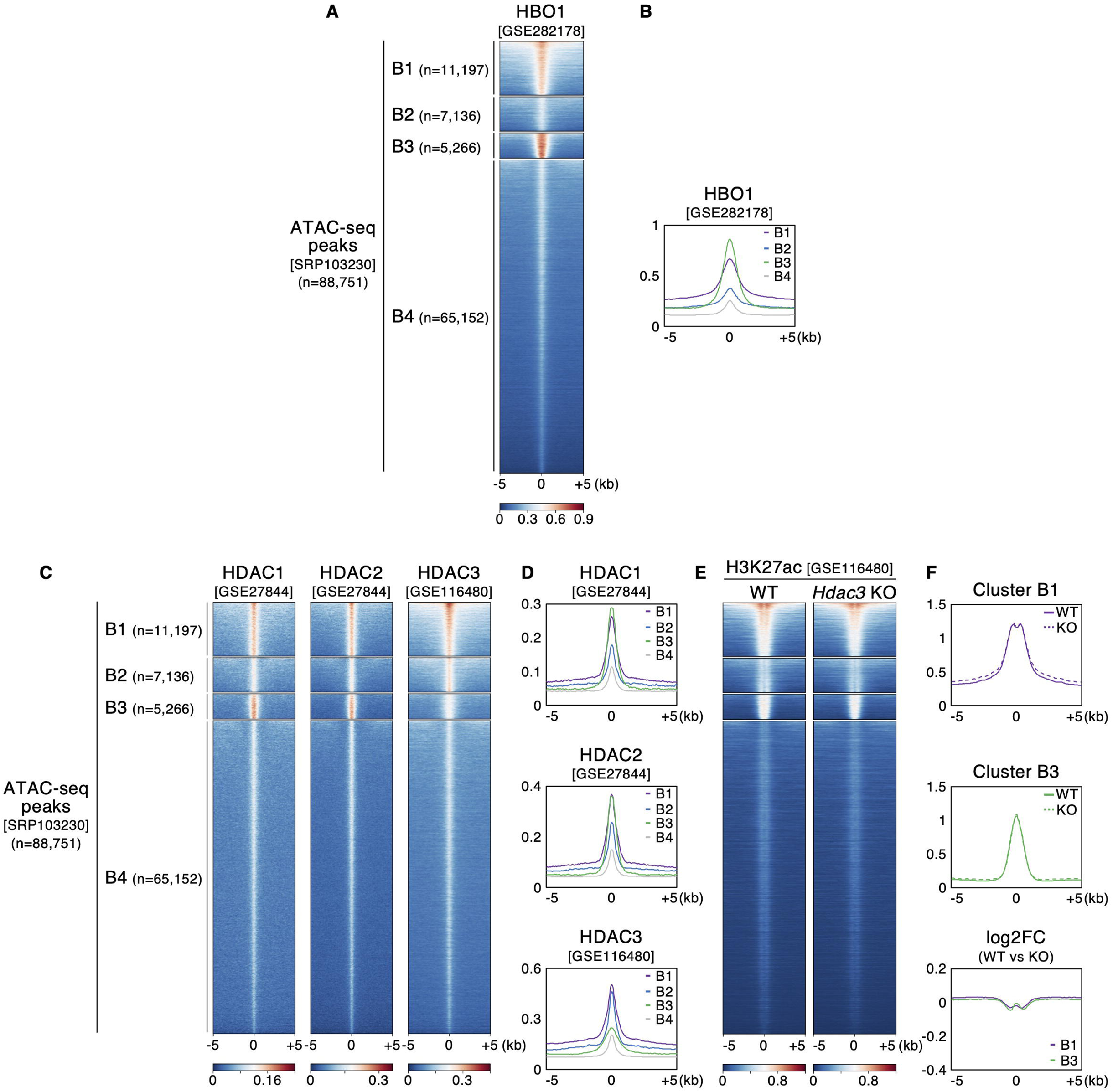
Genome binding of putative histone lactylation regulators in mESCs. (A-D) Heatmaps (A and C) and aggregation plots (B and D) of ChIP-seq signals of HBO1 (A and B) and HDAC1-3 (C and D) at Clusters B1–B4. (C) Heatmaps of H3K27ac signals at Clusters B1–B4 in wild-type (WT) and *Hdac3* knockout (KO) mESCs. (D) Aggregation plots of H3K27ac levels and log2FC values at Clusters B1 and B3 in WT and *Hdac3* KO mESCs.

**Table S1. Public omics datasets used in this study**

**Table S2. Hit lists of TF motif analysis (top 10)**

**Table S3. Hit lists of KEGG pathway analysis (top 10, FDR<0.05)**

## Notes

### Competing Interest Statement

The authors have declared no competing interest.

### Summary of Updates

Figures 5 and S3 added; Figure S2 updated; title, abstract, and main text modified.

